# Parp3 promotes long-range end-joining in murine cells

**DOI:** 10.1101/255281

**Authors:** Jacob V. Layer, J. Patrick Cleary, Alexander J. Brown, Kristen E. Stevenson, Sara N. Morrow, Alexandria Van Scoyk, Rafael B. Blasco, Elif Karaca, Fei-Long Meng, Richard L. Frock, Trevor Tivey, Sunhee Kim, Hailey Fuchs, Roberto Chiarle, Frederick W. Alt, Steven A. Roberts, David M. Weinstock, Tovah A. Day

**Affiliations:** Department of Medical Oncology, Dana-Farber Cancer Institute, Boston, MA, 02215, USA; School of Molecular Biosciences, Washington State University, P100 Dairy Road, Pullman, WA 99164, USA; Department of Biostatistics and Computational Biology, Dana-Farber Cancer Institute, 450 Brookline Avenue, Boston, Massachusetts 02215, USA; Department of Pathology, Children's Hospital, Harvard Medical School, Boston, MA, USA; Howard Hughes Medical Institute, Program in Cellular and Molecular Medicine, Children's Hospital Boston, Department of Genetics, Harvard Medical School, Boston, Massachusetts, 02215, USA; Department of Molecular Biotechnology and Health Sciences, University of Turin, Turin, Italy; Broad Institute of MIT and Harvard University, Cambridge, Massachusetts, 02142, USA.

## Abstract

Chromosomal rearrangements, including translocations, are early and essential events in the formation of many tumors. Previous studies that defined the genetic requirements for rearrangement formation have identified differences between murine and human cells, most notably in the role of classical‐ and alternative-nonhomologous end joining factors (NHEJ). We reported that poly(ADP)ribose polymerase 3 (PARP3) promotes chromosomal rearrangements induced by endonucleases in multiple human cell types. In contrast to c-NHEJ factors, we show here that Parp3 also promotes rearrangements in murine cells, including translocations in murine embryonic stem cells (mESCs), class switch recombination in primary B cells and inversions in tail fibroblasts that generate *Eml4-Alk* fusions. In mESCs, Parp3-deficient cells had shorter deletion lengths at translocation junctions. This was corroborated using next-generation sequencing of *Eml4-Alk* junctions in tail fibroblasts and is consistent with a role for Parp3 in promoting the processing of DNA double-strand breaks. We confirmed a previous report that Parp1 also promotes rearrangement formation. In contrast with Parp3, rearrangement junctions in the absence of Parp1 had longer deletion lengths, suggesting Parp1 may suppress DSB processing. Together, these data indicate that Parp3 and Parp1 promote rearrangements with distinct phenotypes.

---

Chromosomal rearrangements are critical events in the pathogenesis of malignant and nonmalignant disorders (1-3). These aberrant events drive malignant transformation and congenital disorders, including deafness, schizophrenia, and infertility. Many efforts to elucidate the genetic basis of rearrangement formation have relied upon experiments in mouse cells: studies in murine embryonic stem cells (mESCs) identified a cohort of genetic factors that promote or suppress rearrangements (4-8). In aggregate, these studies suggest that chromosomal rearrangements form by a non-canonical or alternative non-homologous end joining pathway (alt-NHEJ). However, a subsequent report demonstrated that in human cells, rearrangement between endonuclease-induced double-strand breaks (DSBs) depends on classical NHEJ (c-NHEJ) (9). The disparate results suggest that the genetic basis for rearrangements differs significantly between human and murine cells.

We recently reported that PARP3, a member of the ADP-Ribose Polymerase family of enzymes, promotes chromosomal rearrangement formation in human cells (10). We showed that PARP3 regulates G quadruplex (G4) DNA in response to DNA damage. Chemical stabilization of G4 DNA in *PARP3*^-/-^ cells led to widespread DSBs. This suggested a model in which PARP3 suppresses G4 DNA, which allows for processing of DSB ends to intermediates that participate in rearrangements in human cells. Here we investigated the role of Parp3 within murine cells using a range of cell types and approaches to quantify frequency and characterize junction phenotypes.

## Parp3 promotes targeted translocations in murine embryonic stem cells

First, we tested the effects of Parp3 depletion using murine embryonic stem cells (mESC) harboring the pCr15 reporter (4). Introduction of the I-*Sce*I endonuclease into these cells leads to concurrent DSBs on chromosomes 14 and 17. Translocation between these DSBs generates a functional neomycin resistance gene on der(17). Knockdown of Parp3 reduced the frequency of targeted translocations induced by I-*Sce*I by approximately 80% compared to a control siRNA (Fig. 1A-C). This is approximately the same extent of reduction previously reported after knockdown of the DSB end-resection factor CtIP (7). PARP3 depletion did not affect I-*Sce*I protein expression, cleavage by I-SceI, or colony plating efficiency (Fig. 1C-E). *PARP3* re-expression after suppression with a UTR-directed siRNA rescued the siRNA effect on rearrangement frequency (Fig. 1F). As previously reported (4), cells lacking the c-NHEJ factor Ku70 had approximately 2-fold higher rearrangement frequency compared to wild-type pCr15 cells (Fig. 1B). PARP3 knockdown also reduced rearrangement frequency in *Ku70*^-/-^ pCr15 cells (Fig. 1B), indicating that PARP3 promotes rearrangements both in the presence and absence of *Ku70*.

**Figure 1.**
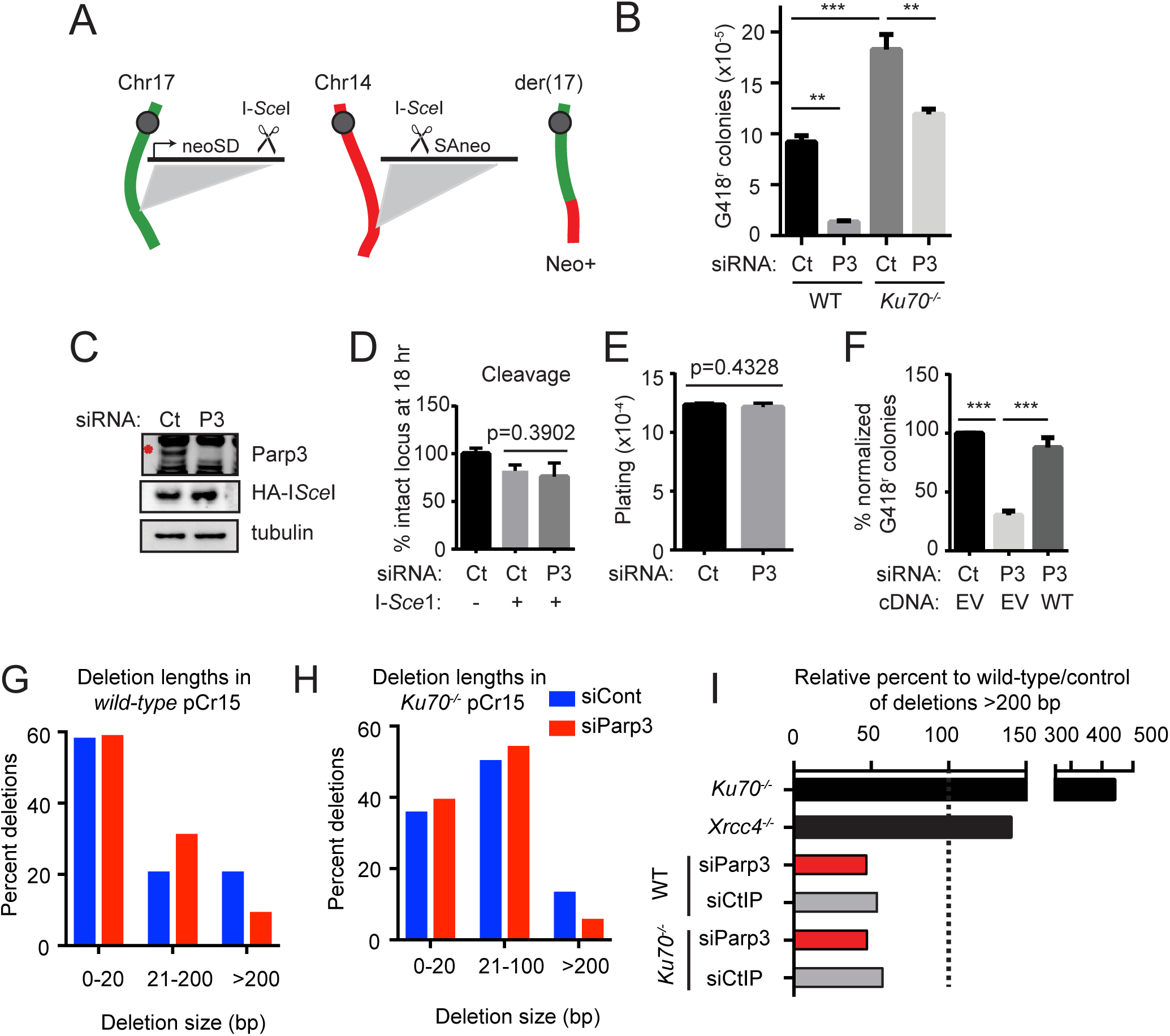
Parp3 promotes translocations in mESCs. (*A*) Schematic of targeted translocation assay in mESCs (4). (*B*) Absolute frequency of translocations in wild-type (WT) and *Ku70*^-/-^ pCr15 mESCs transfected with siRNA against Parp3 (P3) or non-targeting control (Ct). *P* values calculated using a two-way ANOVA with Tukey’s correction. (*C*) Immunoblots in pCr15 cells. Red asterisk, PARP3. Representative experiment of 3 biological replicates. (*D*,*E*) Cleavage by I-*Sce*I measured by quantitative PCR across the targeted site (*D*) and plating efficiency of mESCs (*E*) transfected with siRNA against *Parp3* (P3) or non-targeting control (Ct). *P* values calculated using unpaired Student’s *t* test. (*F*) Frequency of translocations with siParp3 targeting the 5’UTR and re-expression of *Parp3* transcript by transfection of pCAGGS containing either empty vector (EV) or PARP3 (WT). *P* values calculated using two-way ANOVA with Tukey’s correction. (*G*,*H*) Length of deletions at translocation junctions from wild-type (*G*) and *Ku70*^-/-^ (*H*) pCr15 cells. *P* = 0.0055 for siCont vs. siParp3 deletion lengths in wild-type and *p* = 0.0119 for siCont vs. siParp3 deletion lengths in *Ku70*^-/-^. *P*-values were calculated using an unpaired Student’s *t*-test. (*I*) Fold-change in deletions larger than 200 bp reported in pCr15 cells with the indicated genetic perturbations compiled from (4, 6, 7) and this study. Knockout cells are relative to wild-type controls in individual experiments. siRNA knockdowns are relative to siRNA controls in individual experiments. \**P*<0.05, \*\**P*<0.01, \*\*\**P*<0.001.

We examined the characteristics of the rearrangement junctions to elucidate Parp3-dependent mechanisms involved in rearrangement formation. A central challenge of this analysis is to interpret Parp3-dependent changes in rearrangement junctions in the context of a Parp3-dependent reduction in overall rearrangement frequency (Fig. 1B). Therefore, throughout our analysis, we have referred to junction phenotypes in ‘residual’ events. Among the residual translocations in Parp3-depleted cells, there was a notable reduction in junctions with longer deletions (*i.e*., >200 bp) (Fig. 1G,H). This resulted in a statistically significant reduction in mean length of deletions at rearrangement junctions in both wild-type pCr15 cells and *Ku70*^-/-^ pCr15 cells (Fig. 1G,H, Dataset S1). This suggests that Parp3 plays a role in promoting long deletions at rearrangement junctions and that role is Ku70-independent.

We compared the phenotypes reported from previous studies using the pCr15 line to our data (4, 6, 7). In these studies, knockout of the c-NHEJ factors *Ku70* or *Xrcc4* increased the proportion of junctions with >200 bp deletions in residual translocations (Fig. 1I, Dataset S1). In contrast, knockdown of Parp3 led to a similar reduction in the proportion of junctions with >200 bp deletions as reported after knockdown of CtIP (Fig. 1I). We did not find any significant effects from Parp3 knockdown on the proportion of events with insertions or on the usage of microhomology (*i.e*. 1-10 bp stretches of homology) at translocation junctions (Fig. S1A,B, Table 1).

**Table 1.** Repair Statistics in mESCs. Microhomology, deletions, and insertions at I-*Sce*I-mediated translocation junctions in WT and *Ku70*^-/-^ mESCs.

|  | Microhomology |  | Deletions |  |  | Clones with insertion |  | Total clones |
| --- | --- | --- | --- | --- | --- | --- | --- | --- |
| Treatment | Mean (bp) ± SEM | *p*-value | Median | *p*-value | # of clones | Total | % |  |
| Wild-type |  |  |  |  |  |  |  |  |
| siCont | 2.445 ± 0.14 | 0.1172 | 19 | 0.0055 | 149 | 34 | 22 | 153 |
| siParp3 | 2.767 ± 0.1487 |  | 19 |  | 137 | 34 | 25 | 137 |
| Ku70<sup>-/-</sup> |  |  |  |  |  |  |  |  |
| siCont | 3.082 ± 0.1563 | 0.2089 | 36 | 0.0119 | 111 | 27 | 24 | 112 |
| siParp3 | 2.794 ± 0.1644 |  | 27 |  | 101 | 35 | 34 | 102 |

## Parp3 promotes class switch recombination

To explore the *in vivo* phenotypes of *Parp3*-deficiency, we used CRISPR/Cas9 mutagenesis to establish *Parp3*^-/-^ mice. We deleted a 492 bp region containing two out of three Parp3 catalytic residues in mESCs (Fig. S1C-E). Cells with the deletion exhibited complete loss of Parp3 expression (Fig. S1F,G) and *Parp3*^-/-^ mice were established from these mESCs in the 129s background. As previously reported in a separate knockout mouse (11), *Parp3*^-/-^ mice were born in expected Mendelian ratios and had no apparent gross phenotypes.

Based on our observation that *Parp3* promotes targeted rearrangements in mESCs, we hypothesized that Parp3 would also promote class switch recombination (CSR), a physiological form of intrachromosomal rearrangement. However, a recent study reported that Parp3 loss increased CSR frequency (12), which was believed to involve increased occupancy at switch regions by activation-induced deaminase (AID) in the absence of Parp3. Therefore, we first sought to establish whether Parp3 influences AID occupancy at switch regions Sμ-1 and Sμ-2) in our knockout mouse. We performed chromatin immunoprecipitation (ChIP) for AID in wild-type and *Parp3*^-/-^ primary B cells and quantitative PCR (qPCR) for Sμ-1 and Sμ-2. As expected, ChIP-qPCR demonstrated marked enrichment of Sμ-1 and Sμ-2 sequences in wild-type B cells compared to B cells from *Aid*-knockout mice. Unlike the previous report, we did not find any significant differences in AID occupancy at Sμ-1 and Sμ-2 between *Parp3*^-/-^ B-cells and B-cells from wild-type control littermates (Fig. 2A,B).

**Figure 2.**
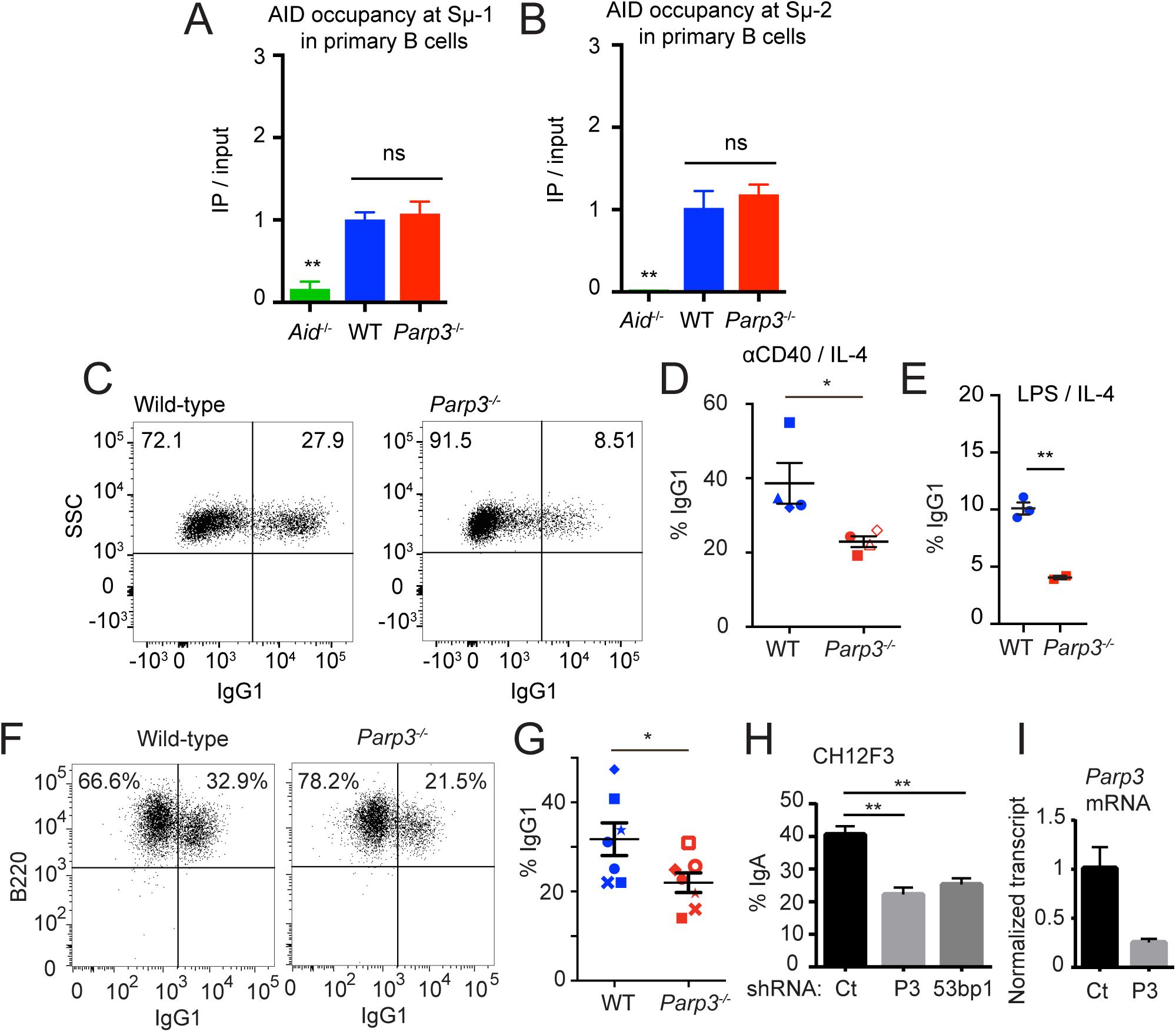
Parp3 promotes CSR in murine B cells. (*A*,*B*) AID occupancy at Sμ1 (*A*) and Sμ2 (*B*) measured by chromatin immunoprecipitation (ChIP). *P* values were calculated using a oneway ANOVA with Tukey’s correction. (*C-E*) *Ex vivo* CSR to IgG1 in primary B cells from WT-or *Parp3*^-/-^ mice following stimulation with aCD40/IL-4 (*C,D*) or LPS/IL-4 (*E*). Data are presented as mean ± SE. *P* values were calculated using unpaired Student’s *t*-test. (*F*,*G*) *Ex vivo* CSR to IgG1 in primary B cells from *Rag2*-deficient blastocyst complementation with WT‐ or *Parp3*^-/-^-deficient mESCs. A representative experiment is shown and data are presented as mean ± SE from 7 different experiments, with each paired experiment indicated by a different symbol. (*H*,*I*) *In vitro* CSR to IgA (*H*) and quantification of *Parp3* transcript (*I*) in CH12F3 cells transduced with shRNA targeting *Parp3*, *53bpl*, or control (Ct). *P* values calculated using a one-way ANOVA with Dunnett’s correction. \**P* < 0.05, \*\**P* < 0.01, \*\*\**P* < 0.001.

The absence of an effect on recruitment within our model allows us to directly investigate the contribution of Parp3 to recombination downstream of AID recruitment. We confirmed that *Parp3*^-/-^ mice have no defects in the early stages of B-cell maturation, which precede CSR (Fig. S2). *Parp3*^-/-^ B cells underwent CSR to IgG1 ~40-50% less efficiently than wild-type cells upon *in vitro* stimulation with either αCD40/IL-4 or LPS/IL-4 (Fig. 2C-E). Despite the reduced frequency of CSR, *Parp3*^-/-^ B cells had similar or increased AID expression, Sμ-1 and Sμ-2 transcript levels or *in vitro* proliferation following stimulation compared to wild-type B cells (Fig. S3A-D). Analysis of switch junction sequences from stimulated B cells revealed that there was no significant Parp3-dependent change in insertions or microhomology usage (Fig. S3E).

To confirm these findings with a second approach, we performed *Rag2*-deficient blastocyst complementation (13) to reconstitute the lymphoid compartment with our *Parp3*^-/-^ mESCs. After *in vitro* stimulation with αCD40/IL-4, *Parp3*^-/-^ B cells isolated from complemented *RagI*^-/-^ mice had ~35% reduced CSR to IgG1 compared to wild-type B cells (Fig. 2F,G). As a third approach, we used the CH12F3 murine B cell line, which undergoes CSR from IgM to IgA upon stimulation with αCD40, IL-4, and TGF-β (14). Compared to wild-type cells, shRNA-mediated knockdown of *Parp3* reduced CSR to IgA by ~50% compared to cells with control shRNA (Fig. 2H,I). This reduction was similar in effect to shRNA-mediated knockdown of *53bp1* (Fig. 2H), a factor known to promote CSR (15, 16).

## Loss of Parp3 reduces *Eml4-Alk* inversions

To study the phenotype of rearrangement junctions in murine cells in more depth, we examined inversions between the *Eml4* and *Alk* loci on mouse chromosome 17 (Fig. 3A). We chose *Eml4-Alk* because it is a known driver of non-small cell lung cancer in both mice and humans (17, 18). In addition, the relatively high frequency of rearrangements in this system (19) coupled to high-throughput analysis of amplicon deep sequencing allowed us to compare *Eml4-Alk* rearrangement junctions (*i.e*., distal repair events) to non-rearrangement repair events at the *Alk* locus (*i.e*., proximal repair events). The murine *Alk* locus is predicted to contain abundant G4 DNA structures (Fig. S3F). Thus, we hypothesized that Parp3 deletion would affect chromosomal rearrangements involving this locus.

**Figure 3.**
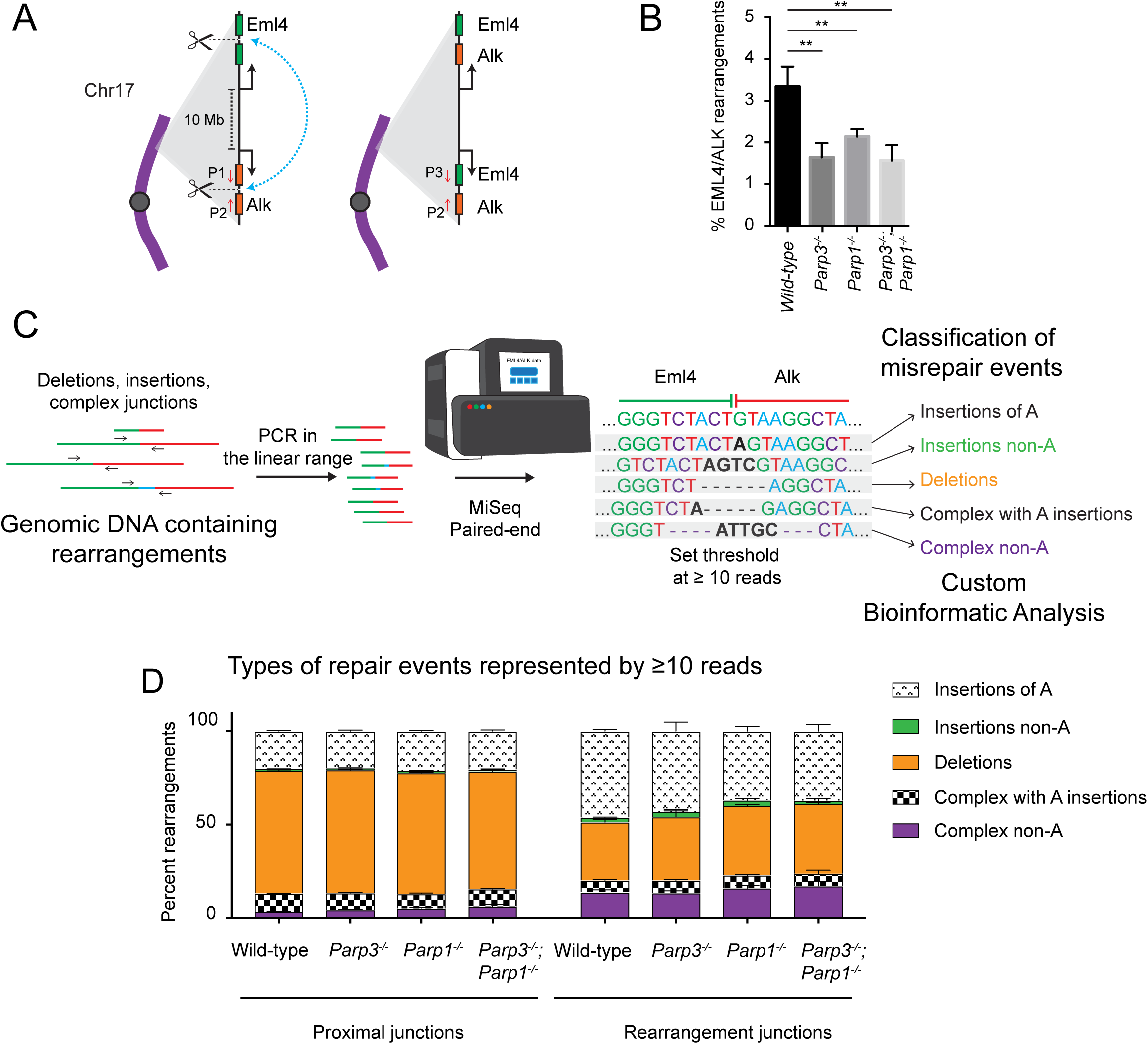
High throughput analysis of amplicon deep sequencing. (*A*) Schematic of *Eml4-Alk* rearrangements on mouse chromosome 17. CRISPR/Cas9 cleavage in the *Eml4* and *Alk* loci can lead to a 10 Mb inversion that creates an oncogenic fusion. Scissors, Cas9-mediated DSB. P1, P2, P3 are PCR primers. P1 and P2 amplify proximal repair. P3 and P2 amplify rearrangements. (*B*) Frequency of *Eml4-Alk* rearrangements in murine tail fibroblasts measured by droplet digital PCR 5 days after adenovirus-mediated expression of CRISPR/Cas9. *P* values were calculated using a one-way ANOVA with Dunnett’s correction. (*C*) Schematic of sequencing and analysis pipeline for repair amplicons. PCR in the linear range generates amplicons from proximal repair or rearrangements. MiSeq paired-end sequencing is followed by trimming and mapping of reads to reference sequences. Differences from the reference sequence are classified as deletions, insertions, or complex repair events. Insertions or complex repair events containing a single ‘A’ nucleotide insertion are designated separately as they are likely CRISPR/Cas9-mediated. (*D*) Distributions of types of repair events by represented by misrepair sequences with ≥10 reads. Data are presented as mean ± S.E. of *n*=3. \**P* < 0.05, \*\**P* < 0.01, \*\*\**P* < 0.001.

In cultures of primary murine tail fibroblasts, we used an established adenoviral approach to express CRISPR/Cas9 with gRNA targeting intron 13 of *Eml4* and intron 19 of *Alk* (Fig. 3A) (19, 20). We performed droplet digital PCR (ddPCR) to measure the frequencies of rearrangements and observed that rearrangement frequency was reduced ~50% in *Parp3*^-/-^ tail fibroblasts (Fig. 3B) without an appreciable difference in adenoviral transduction efficiency (Fig. S3G).

## Differences in repair junction phenotypes between *Parp3*^-/-^ and *Parp1*^-/-^ cells

High-throughput analysis of amplicon deep sequencing can facilitate the rapid interrogation of repair phenotypes (21-23). We used this approach to address an aspect of the long-standing question of PARP enzyme redundancy in the context of DNA repair. Parp1 and Parp3 share ~60% homology in their catalytic and WGR domains (24), modify overlapping and distinct targets in DNA repair (11, 25) and both promote chromosomal rearrangements in murine cells (Fig 1B and (8)). Together, these findings suggest that they could act by a common mechanism. To examine the relationship between Parp3 and Parp1 in rearrangement formation, we first compared *Eml4-Alk* rearrangement frequencies. *Parp1*^-/-^ tail fibroblasts exhibited ~50% reduced rearrangement frequency (Fig. 3B), similar to *Parp3*^-/-^ tail fibroblasts. Tail fibroblasts lacking both *Parp3*^-/-^ and *Parpl*^-/-^ also had ~50% reduced rearrangement frequency (Fig. 3B), suggesting the enzymes are epistatic with respect to this phenotype.

Next, we used PCR in the linear range (Fig. S3H) to amplify ~250 bp regions surrounding either the *Eml4-Alk* rearrangement junction or the *Alk* CRISPR/Cas9 cut site (Fig. 3A). The amplicons, including controls from cells without CRISPR/Cas9 cutting, were deep sequenced in multiplexed, paired-end MiSeq reactions. Any ‘misrepaired’ sequences found only in the uncut controls were assumed to be sequencing artifacts and excluded from our analysis. Similar to recent studies, our analysis cannot distinguish uncut loci from error-free repair (21, 22). Therefore, we only considered sequences with insertions or deletions to be unambiguous products of end joining. We used a custom bioinformatic analysis to categorize all misrepaired sequences represented by ≥10 reads as deletions, insertions or complex repair events (deletion combined with insertion) (Fig. 3C) (21, 22). We set our threshold at ≥10 reads for a unique junction to reduce PCR artifacts and increase the likelihood that repair events would be reproducibly observed across samples. Three biological replicates were performed for each of the four genotypes for both *Eml4-Alk* rearrangement and proximal repair at the *Alk* locus. We determined the relative frequencies of different repair events by dividing by the total number of reads (for rearrangements) or by the total number of misrepaired reads (proximal repair) (Fig. 3D, S3I). In addition, we analyzed all misrepaired sequences represented by ≥1 reads rather than the threshold of ≥10 reads and it yielded very similar distributions (Fig. S4A).

We first noted an unexpectedly high proportion of both proximal repair junctions and rearrangements were classified as insertions (~25% and ~50% respectively Fig. 3D). Of these insertions, the large majority consisted of a single ‘A’ base pair (Fig. 3D and Table S1), a common repair outcome for a CRISPR/Cas9 DSBs (26). Among the complex junctions, ~35-75% consisted of 1 bp insertions of ‘A’ concurrent with deletions (Fig. 3D). For the purposes of our initial analysis, we retained these events within the insertion and complex categories. We compared the distributions of repair events found at rearrangement junctions versus proximal junctions. To do so, we constructed a multinomial logistic regression model with a random effect for repeated measures using type of junction as the dependent variable (Table S2). Due to the large number of reads in the analysis (Fig. S3I), most comparisons were significant (*p* < 0. 0001), so we only considered those with an odds ratio (OR) ≥1.25 or ≤ 0.75 to be of interest. Rearrangements in each genetic background (wild-type, *Parp3*^-/-^, *ParpV*^-/-^ and *Parp3*^-/-^; *Parp1*^-/-^ were less likely to have deletions and more likely to have insertions relative to proximal repair in the same genetic background (*p* < 0.0001, ORs 4.93, 4.05, 3.13, and 3.05 respectively, Fig. 3D, Table S2). As the vast majority of these insertions are likely CRISPR/Cas9-mediated, we evaluated the dataset neglecting all events containing insertions of single ‘A’ nucleotides and found similar patterns for all genotypes (*p* < 0.0001, ORs 6.03, 4.72, 4.47, and 2.61 respectively, Fig. S4B, Table S3).

Using the complete dataset, we compared distributions of rearrangement repair events between genotypes. Rearrangements in *Parp3*^-/-^ or *Parp1*^-/-^ cells were more likely to have deletions than those in wild-type cells (OR = 1.25, OR = 1.48 respectively, Table S2, Figure 3D). Cells lacking both enzymes exhibited a pattern similar to either single knockout (OR = 1.52, Table S2, Figure 3D). In addition, rearrangements in both *Parp1*^-/-^ and *Parp3*^-/-^;*Parp1*^-/-^ cells, were less likely to have insertions than those in wild-type cells (OR = 0.7, OR = 0.64 respectively, Table S2). In comparisons across genotypes for proximal repair, the odds ratios were < 1.25 or > 0.75, indicating that the differences between distributions of repair events during proximal repair of the *Alk* locus were less dramatic than in *Eml4-Alk* rearrangements (Fig. 3D). In the analysis of the dataset without single A insertions, rearrangements in *Parp3*^-/-^;*Parp1*^-/-^ cells remained less likely to have insertions than wild-type cells (OR = 0.47, Fig. S4B, Table S3). However, in this dataset, the *Parp3*‐ and *Parp1*-dependent effects were extinguished (Fig. S4B, Table S3), suggesting that they were CRISPR/Cas9-mediated effects. Taken together, during rearrangement formation, combined loss of *Parp3* and *Parp1* decreases insertions

Next, we examined mean lengths of deletions, microhomologies, and insertions (Table S4) in the residual rearrangement junctions. In wild-type cells, we observed that the mean deletion length in rearrangement junctions was significantly longer than in proximal junctions (Fig. 4A, Table S4) consistent with the finding that rearrangements in murine cells involve more extensive processing of DNA breaks (4, 27). We observed the same pattern in *Parp1*^-/-^ and *Parp3*^-/-^;*Parp1*^-/-^ cells (Figure 4A, Table S4). In contrast, in *Parp3*^-/-^ cells, the mean deletion lengths for rearrangements and proximal repair were nearly the same (Figure 4A, Table S4).

**Figure 4.**
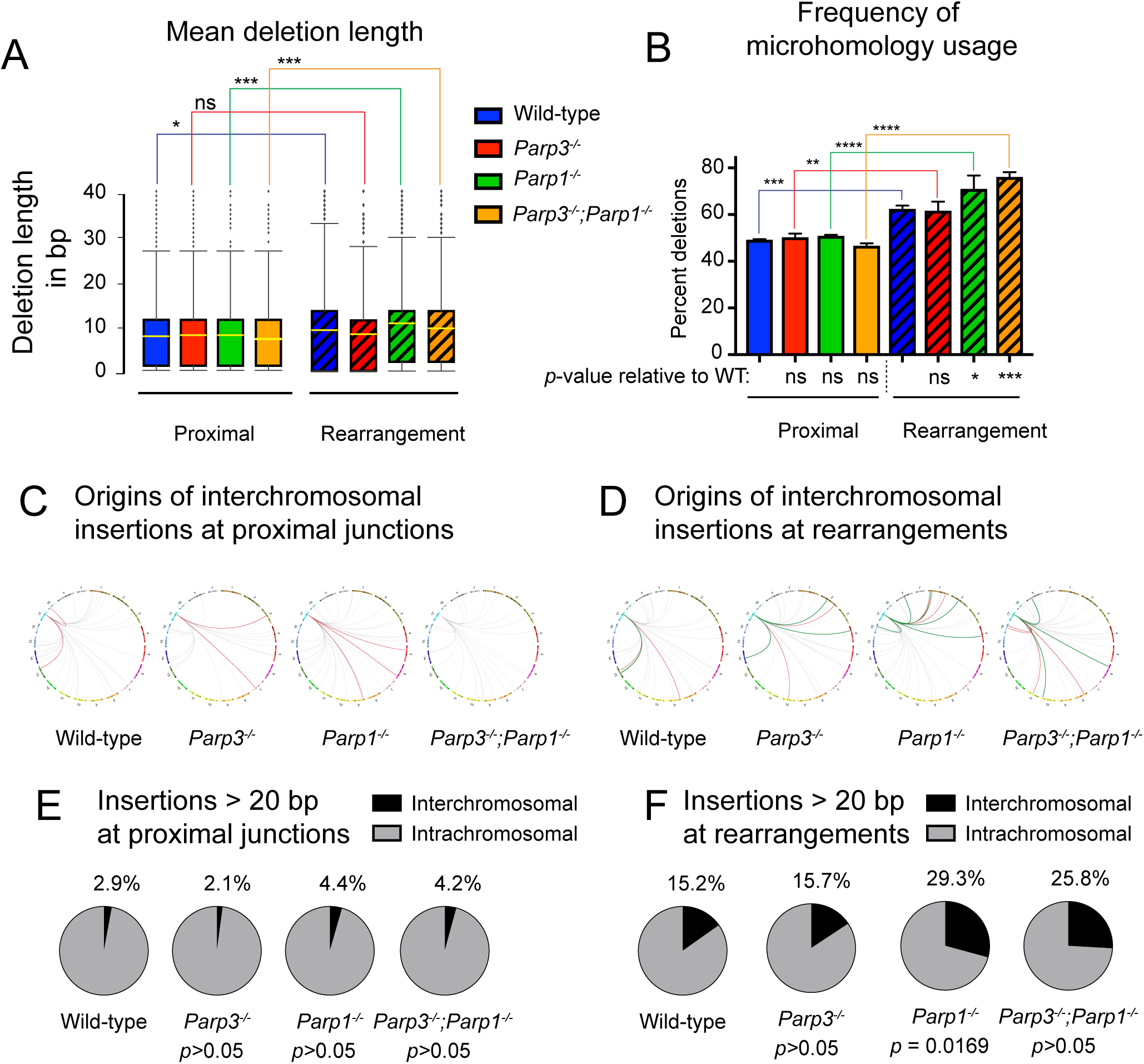
Repair phenotypes in murine tail fibroblasts. (*A*) Distribution of deletion lengths in proximal repair and at rearrangements. Box represents the quartile range, solid lines extend to minimum and 99^th^ percentile, diamonds above maximum are outliers, yellow line is mean deletion length. (*B*) Percent of deletions that exhibit microhomology. *P* values were calculated using a two-way ANOVA with Tukey’s correction. *P* values beneath the x-axis compare each genotype to wild-type value within that repair location (i.e. proximal or rearrangement). (*C*,*D*) Genomic distributions of the origins of templated interchromosomal insertions ≥20 bp in length in proximal repair (*C*) and rearrangements (*D*). (*E*,*F*) Proportion of inter‐ and intra-chromosomal templated insertions ≥20 bp in length in proximal repair (*E*) and at rearrangements (*F*). *P* values calculated using a two-sided Chi Square. \**P* < 0.05, \*\**P* < 0.01, \*\*\**P* < 0.001. ns, not significant.

Based on our previous report (10), we hypothesized that Parp3 promotes the processing of DNA DSBs during rearrangement in murine cells, which results in repair that involves deletions of end sequence. As expected, rearrangements in *Parp3*^-/-^ cells had a shorter mean deletion length than those in wild-type cells. In contrast, rearrangements in *Parp1*^-/-^ cells had longer mean deletion length than wild-type cells. These data suggest opposing trends though neither difference achieved statistical significance (*p* = 0.1565 and *p* = 0.0525 respectively, Fig. 4A, Table S4). In support of these trends, rearrangements in *Parp1*^-/-^ cells had significantly longer deletions than those in *Parp3*^-/-^ cells (*p* = 0.0001, Fig. 4A). In addition, rearrangements in *Parp3*^-/-^;*Parp1*^-/-^ cells had significantly shorter deletions than those in *Parp1*^-/-^ cells (*p* = 0.0202, Fig. 4A). Taken together, these data suggest that during rearrangement formation, *Parp3* promotes and *Parp1* suppresses DSB processing in murine fibroblasts. Within the proximal repair junctions, we did not find any statistically significant differences in mean deletion lengths between genotypes (Fig. 4A, Table S4).

During DSB repair, deletions can uncover microhomologies that may be preferentially utilized to facilitate repair by alt-NHEJ (28). Reports in the literature indicate that rearrangements in murine cells occur predominantly by alt-NHEJ (4-8). A comparison of the flanking sequences indicated comparable opportunities for microhomology usage during proximal repair or during *Eml4/Alk* rearrangement (Fig. S4C). Yet, rearrangements exhibited significantly increased frequencies of microhomology usage compared to proximal repair in wild-type, *Parp1*^-/-^, *Parp3*^-/-^, and *Parp3*^-/-^;*Parp1*^-/-^ cells (Fig. 4B). Mean length of microhomology was also significantly increased at rearrangements compared to proximal repair for all 4 genotypes (Fig. S4D).

Among rearrangements, there were no significant *Parp3*-dependent differences in either proportion of repair events with microhomology or mean length of microhomology (Figs. 4B, S4D), consistent with our findings in pCr15 cells and *Parp3*^-/-^ B cells (Figs. S1A and S3E). In contrast, *Parp1*^-/-^ and *Parp3*^-/-^;*Parp1*^-/-^ cells had significantly increased frequencies of microhomology usage at rearrangements compared to wild-type cells (Fig. 4B). In addition, rearrangements in *Parp3*^-/-^;*Parp1*^-/-^ cells exhibited significantly longer mean microhomology usage than those in wild-type cells (Fig. S4D). No significant differences in microhomology were observed in proximal repair (Figs. 4B and S4D).

To evaluate mean insertion lengths during DSB repair, we included both simple insertions and insertions occurring with a deletion (complex events). Loss of Parp1 led to a significant increase in mean insertion length (Fig. S4E, Table S4) in both proximal repair and rearrangements. In double knockout cells, mean insertion lengths were also increased although the increase in proximal repair did not achieve statistical significance (Fig. S4E, Table S4). No additional Parp3-dependent effect was observed, indicating the increases in insertion length depend on Parp1. Because CRISPR/Cas9 can lead to a predominance of single nucleotide insertions during repair (26), we also evaluated insertion lengths without these events. When we excluded single ‘A’ insertions, the overall mean insertion lengths were increased, but the Parp1-dependent increase in insertion length remained statistically significant in proximal repair. (Fig. S4E).

Given that each insertion is a unique event, we reasoned that some insertions in our dataset would be represented by fewer than 10 reads. Therefore, to understand the origins of inserted sequences in our system, we combined the three biological replicates for each condition and considered all insertions (both simple insertions and insertions in complex junctions) represented by ≥1 read. We aligned every insertion ≥20 base pairs with genomic sequence and normalized its representation by the number of reads for that sequence. None of the sequences that we interrogated showed significant alignment to the adenoviral genome (29) or Cas9 sequence. Overall, ~99.8% of insertions >20 bp aligned with the mouse genome. Upon manual curation of the remaining 0.2%, we observed single insertions containing sequence from multiple genomic loci; we excluded these events from further analysis.

We divided all aligned insertions from both rearrangement and proximal junctions into sequences that originated from chr.17 or non-chr.17 locations. We noted that insertions from non-chr.17 locations (i.e., interchromosomal insertions) were distributed throughout the genome without any appreciable chromosomal bias (Fig. 4C,D). For all genotypes, the fraction of insertions that involved non-chr.17 sequence was higher in rearrangements than at proximal junctions (Fig. 4E,F). Both *Parp1*^-/-^ and *Parp1*^-/-^;*Parp3*^-/-^ cells had approximately two-fold increases in the proportion of non-chr.17 sequence insertions compared to wild-type cells (Fig. 4E,F; *p* = 0.0169 for rearrangements), suggesting that Parp1 suppresses these events. In conclusion, our data indicate that while loss of either Parp3 or Parp1 leads to a reduction in chromosomal rearrangements, rearrangement junctions in *Parp3*^-/-^ and *Parp1*^-/-^ cells exhibit different phenotypes (Fig. 5).

**Figure 5.**
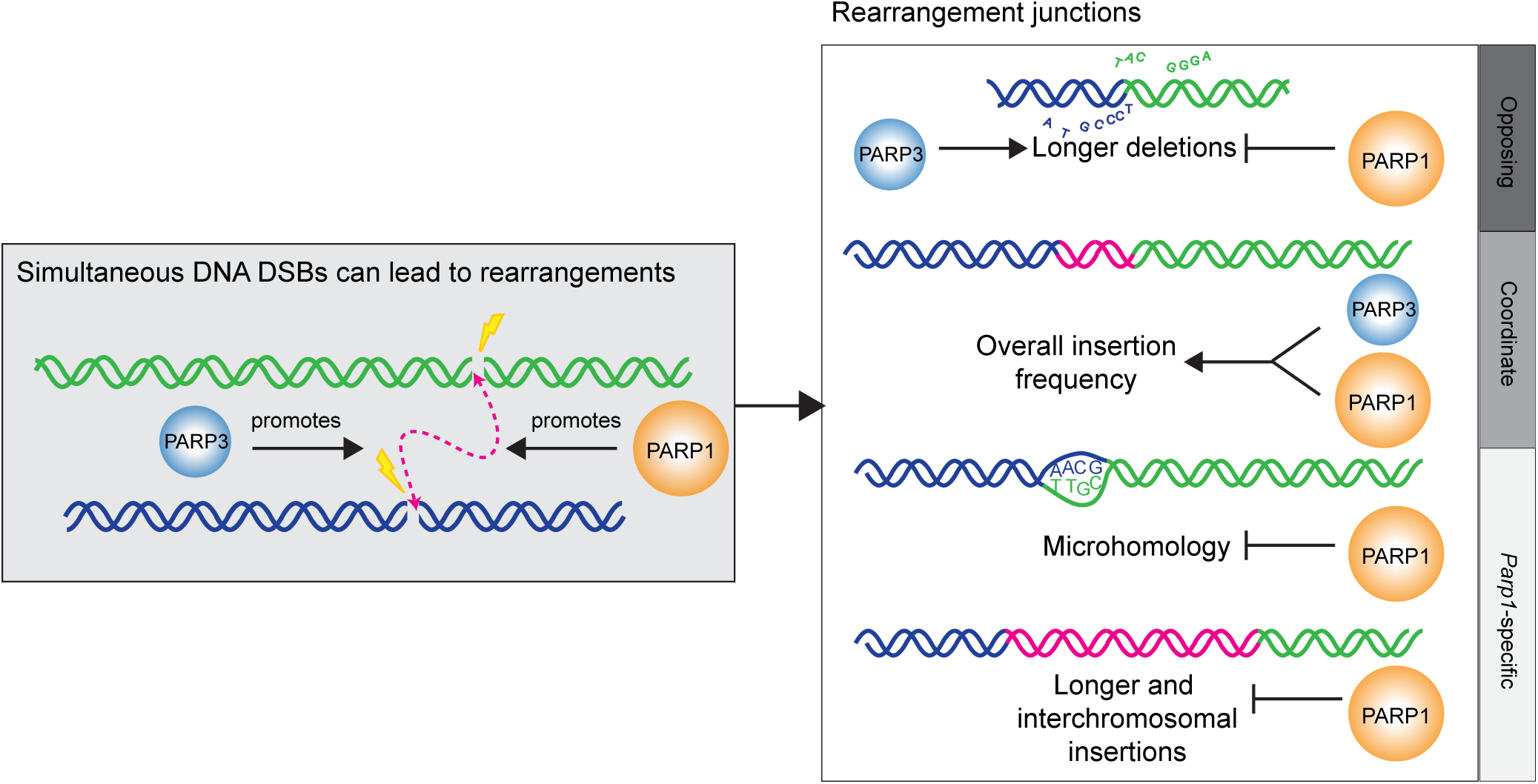
Model of Parp3 and Parp1 participation in rearrangement formation. Both Parp3 and Parp1 promote chromosomal rearrangements. Parp3 promotes longer deletions and Parp1 suppresses them at rearrangements. Parp3 and Parp1 act coordinately to promote insertions at rearrangements. Parp1 uniquely suppresses microhomology use, longer insertions, and interchromosomal insertions at rearrangements.

## Discussion

Expanding upon our previous study in human cells (10), we show here that Parp3 also promotes chromosomal rearrangements in murine cells. We confirm this finding in multiple assays, including I-*Sce*I-mediated translocations in mESCs, class switch recombination in primary B cells, and inversions of the *Eml4/Alk* locus in tail fibroblasts (Figs. 1, 2, and 3). To date, most factors known to mediate chromosomal rearrangements have species-specific roles (4, 7, 9); Parp3 is an exception to this paradigm. Whereas the alt-NHEJ pathway mediates rearrangements in murine cells, c-NHEJ promotes rearrangements in human cells (9). The observation that Parp3 promotes rearrangements in both murine and human cells suggests that Parp3 either 1) acts in both pathways or 2) performs a function upstream of both pathways. Supporting the latter, we previously reported that PARP3 suppresses G quadruplex (G4) DNA to facilitate DSB repair in human cells (10). In murine cells, the *Alk* locus is predicted to contain abundant G4 DNA structures (Fig. S3F), raising the possibility that Parp3 could act upstream of both NHEJ pathways by suppressing G4 DNA in mouse cells as well.

Immunoglobulin switch regions form abundant G4 structures that may be involved in recruitment and oligomerization of AID (30). In our Parp3 knockout, we do not find evidence for Parp3-dependent suppression of AID recruitment, which contrasts with a report by Robert *et al*. (12). We noted that while our *Parp3*^-/-^ cells (both the germ line knockout mouse and the cells used for Rag-deficient blastocyst complementation) are in the 129 background (31), Robert *et al* established their *Parp3*^-/-^ mice in a B6;129 mixed background (12). As significant differences have been noted in antibody class switching between strains of inbred mice (32), this could explain the observed differences.

To deepen our understanding of Parp3 in rearrangement formation, we used high throughput analysis of amplicon deep sequencing. This strategy has enormous applications for rapid, in-depth examination of DNA repair phenotypes (21, 22, 33). The tractability of this technique facilitated comparisons between rearrangement sequences and proximal repair and allowed us to expand our analysis beyond Parp3 to dissect the role of a closely-related enzyme, Parp1. Though Parp3 and Parp1 both promote rearrangements, they appear to do so by distinct mechanisms: Parp1-deficient cells have an increase in longer deletions at rearrangement junctions suggesting that Parp1 suppresses longer deletions. In contrast, rearrangement junctions in Parp3-deficient cells have fewer long deletions (Fig. 5). The same phenotype was observed in mESCs with the pCr15 translocation reporter after Parp3 knockdown (Fig. 1).

While high throughput analysis of deeply sequenced amplicons has great potential, certain aspects of our analysis should be emphasized. Owing to the size of the dataset, even slight differences are likely to achieve statistical significance (Tables S2,3). Therefore, to focus on meaningful differences, it is essential to establish thresholds for analysis. In the present study, we set thresholds to focus on effects that changed likelihood outcomes by ≥25% and reduce artifactual effects. CRISPR/Cas9 has the power to create DNA damage virtually anywhere in the genome, but we observed that it could also drive the predominance of a specific repair event ((26) and Tables S1). As a result, analysis of repair of CRISPR/Cas9 DSBs must account for these potential biases (Figs. 3D and S4B). In addition, use of short PCR amplicons (200-300 bp) coupled to paired end MiSeq confers depth to the assay but limits the window of investigation. In its present iteration, the assay cannot capture deletions or insertions larger than approximately 100-150 bp on either side of the junction. As technology evolves, it may be feasible to use longer amplicons (34) for high-throughput analysis of repair events. Finally, previous studies have reported long templated insertions corresponding to sequences on the DNA plasmid used to express an endonuclease (5, 35, 36). As we do not find any evidence of insertions originating from our adenoviral sequences, adenovirus-mediated delivery may be a successful strategy to avoid such artifactual events.

In conclusion, we find that Parp3 promotes rearrangements in a locus‐ and species-independent manner. We harnessed the power of high throughput analysis of deeply sequenced amplicons to find that Parp3-deficient cells have increased deletion lengths at rearrangement junctions. Though the closely related enzyme Parp1 also promotes rearrangement formation, we uncovered phenotypic differences at residual junctions that support differences in function at DSBs. Our results demonstrate the promising applications of this experimental technique to efficiently evaluate a large number of variables for DNA repair outcomes.

## Methods

### Translocations in pCr15 cells

1 x 10^6^ wild-type or *Ku70*^−/−^ pCr15 cells were transfected with 60 pmol siRNA targeting Parp3 (ON-TARGETplus Mouse Parp3 (L055054-01), Dharmacon) or a non-targeting control (ON-TARGETplus Non-targeting Pool (D-001810-10), Dharmacon) and, for rescue experiments, (ON-TARGETplus Mouse Parp3 (235587) (J-055054-11), Dharmacon). The DNA mix for the first transfection in the rescue experiment included 3 μg pCAGGS PARP3 WT or EV and was performed with Lipofectamine 2000 (Invitrogen). Cells were then cultured for 24 h before nucleofection (Amaxa, Lonza) with 60 pmol of siRNA and 20 μg of either pCAGGS EV or pCAGGS I-*Sce*I. A small fraction of cells were plated after serial dilution and cultured without G418 selection to determine plating efficiency, whereas the majority of cells were plated without dilution and selected in 200 μg/mL G418 after 24 h. Translocation frequency was determined by normalizing the number of *neo*^+^ clones scored after 10 days of G418 selection to plating efficiency.

### Translocation junction analysis

Junction analysis was previously described in (4). Briefly, *neo*^+^ clones were amplified in 96-well plates and lysed by adding 35 μl lysis buffer (10 mM Tris, pH 8, 0.45% (v/v) NP-40, 0.45% (v/v) Tween 20, 100 μg/ml proteinase K) and placing the plates at 55°C for 2 h and 95°C for 5 min. PCR amplification was performed with 2 μL cell lysate per 20 μL reaction using Bioneer Accupower PCR PreMix with the following protocol: 3 min at 94°C and then 40 cycles of 30 sec at 94°C, 1 min at 56°C and 2 min at 72°C. PCR products were treated with ExoSAP-IT (Affymetrix) and sequenced.

### Isolation of murine tail fibroblasts

Isolation of primary murine tail fibroblasts was previously described (19). Briefly, mouse tails were sprayed with 70% ethanol, minced, and incubated overnight in 1.6 mg/mL collagenase type II (Gibco), dissolved in tail fibroblast media. The following day, digested tissue was disrupted by passage through pipettes, cells were centrifuged for 5 minutes at 1200 rpm, and resuspended in tail fibroblast media.

### Droplet Digital PCR

Droplet digital PCR (ddPCR) reactions containing SuperMix (BioRad, 1863024), droplet generator oil, primers and template were mixed and subjected to droplet generation using the QX200 Droplet Generator (BioRad). PCR was performed according to the manufacturer’s instructions and reactions were analyzed on the QX200 Droplet Reader (BioRad). Primer and probes included the TaqMan™ Copy Number Reference Assay, mouse, *Tfrc* (Life Technologies, 4458366) and those found in Supporting Methods.

### Adenovirus Infection

Murine tail fibroblasts were maintained in 0.5% FBS for 24 hours prior to adenovirus infection. Fibroblasts were routinely infected at a multiplicity of infection (MOI) of 50 viral particles per mL. The CRISPR/Cas9 adenovirus with *Eml4*‐ and *Alk*-targeted guides was previously described (19). As a control for adenoviral infection, cells were infected with Ad-Control or Ad-dsRed that were previously described (37).

### Isolation of genomic DNA, PCR and sequencing

Five days after infection with CRISPR/Cas9 adenovirus, genomic DNA (gDNA) was collected from the each population according to the manufacturer’s protocol (Qiagen QIAmp kit). PCR in the linear range was performed using Bioneer Accuprime with 100 ng of gDNA template for each reaction. PCR conditions were 3 minutes at 95°C, followed by cycles of 15 seconds at 95°C, 15 seconds at 55°C, and 15 seconds at 72°C. 27 cycles and 33 cycles were performed for the proximal repair at the *Alk* locus and the *Eml4-Alk* rearrangement respectively. Eight PCR products were pooled for each condition and purified using the Qiagen PCR purification kit. Purified products were visualized on an agarose gel to ensure the presence of a single band. Sequencing on a MiSeq (Illumina) was performed by the CCIB DNA Core Facility at Massachusetts General Hospital (Cambridge, MA) using a MiSeq v2 chemistry 300 cycle kit.

### Amplicon Analysis

PCR amplicons containing DNA repair junctions were analyzed as previously described (21, 22, 33). Briefly, raw reads from paired end sequencing were trimmed to amplicon primer sequences and merged into single reads using Geneious v10.1.3. Junctions with less than 20 base pairs (bp) of reference sequence adjacent to primer sequences were discarded as PCR artifacts. Junctions were trimmed to common starting and ending sequences, mapped to the *Alk* reference sequence or the predicted *Eml4-Alk* blunt-join reference sequence (Genome build GRCm38.p6), and exported as SAM files using Geneious v10.1.3. Using custom Python scripts, the CIGAR strings for each sequenced amplicon were analyzed to classify sequences as exact, deletion, insertion, or complex (containing both deletion and insertion) and to determine the extent of deletions, insertions, and microhomology usage within each sequence.

For the analysis of repair classes and lengths of deletions, insertions, and microhomology, junctions with ≥ 10 reads were analyzed to minimize the effects of PCR artifacts observed in the uncut control. Within each replicate, the normalized percentage of repair class was determined by dividing the number of reads corresponding to deletion, insertion, or complex sequences by the total number of ‘misrepair’ reads. ‘Misrepair’ reads are all reads containing a deletion, insertion, or complex repair but not exact sequences. Deletions are defined as all deletions occurring in deletion and complex sequences. Insertions are defined as all insertions occurring in insertion and complex sequences. Average deletion, insertion, and microhomology lengths were calculated by multiplying the read counts associated with each deletion length, insertion length, or microhomology length, and dividing by the total read counts associated with deletions and complex events (for deletions), insertions and complex events (for insertions), or just deletion events (for microhomology). Potential microhomology usage at the *Alk* and *Eml4-Alk* rearrangements was identified using custom Python scripts.

For analysis of templated insertions, all insertions ≥20bp were considered for further analysis. Any deletions that would eliminate a primer sequence were removed from the dataset. The remaining sequences were submitted to BLAST under default or short sequence parameters against a local *mus musculus* refseq genomic GRCm38.p6 database ftp://ftp.ncbi.nlm.nih.gov/genomes/refseq/vertebrate_mammalian/Mus_musculus/latest_assembly_versions/ and the top hits were exported as XML files for further analysis. Insertions were then classified as inter or intra-chromosomal and for submitted to Circos 0.69-6 to generate Circos plots.

## Acknowledgments

We thank Dipanjan Chowdhury, Andrew Lane and members of the Price and Weinstock laboratories for thoughtful comments. This work was supported by American Cancer Society and Alex Lemonade Stand fellowships (both to T.A.D.), the Claudia Adams Barr Fund for Basic Cancer Research (to T.A.D and D.M.W), a National Science Foundation Graduate Research Fellowship under Grant No. (**DGE1144152**) (to J.V.L), and NIH/NCI R01 CA151898 and R01 CA172387 (both to D.M.W.).

